## Supplementary material for "Intrahost SARS-CoV-2 k-mer identification method (iSKIM) for rapid detection of mutations of concern reveals emergence of global mutation patterns": Figure S1

### Mutations Found in the Alpha Variant (B.1.1.7 lineage)

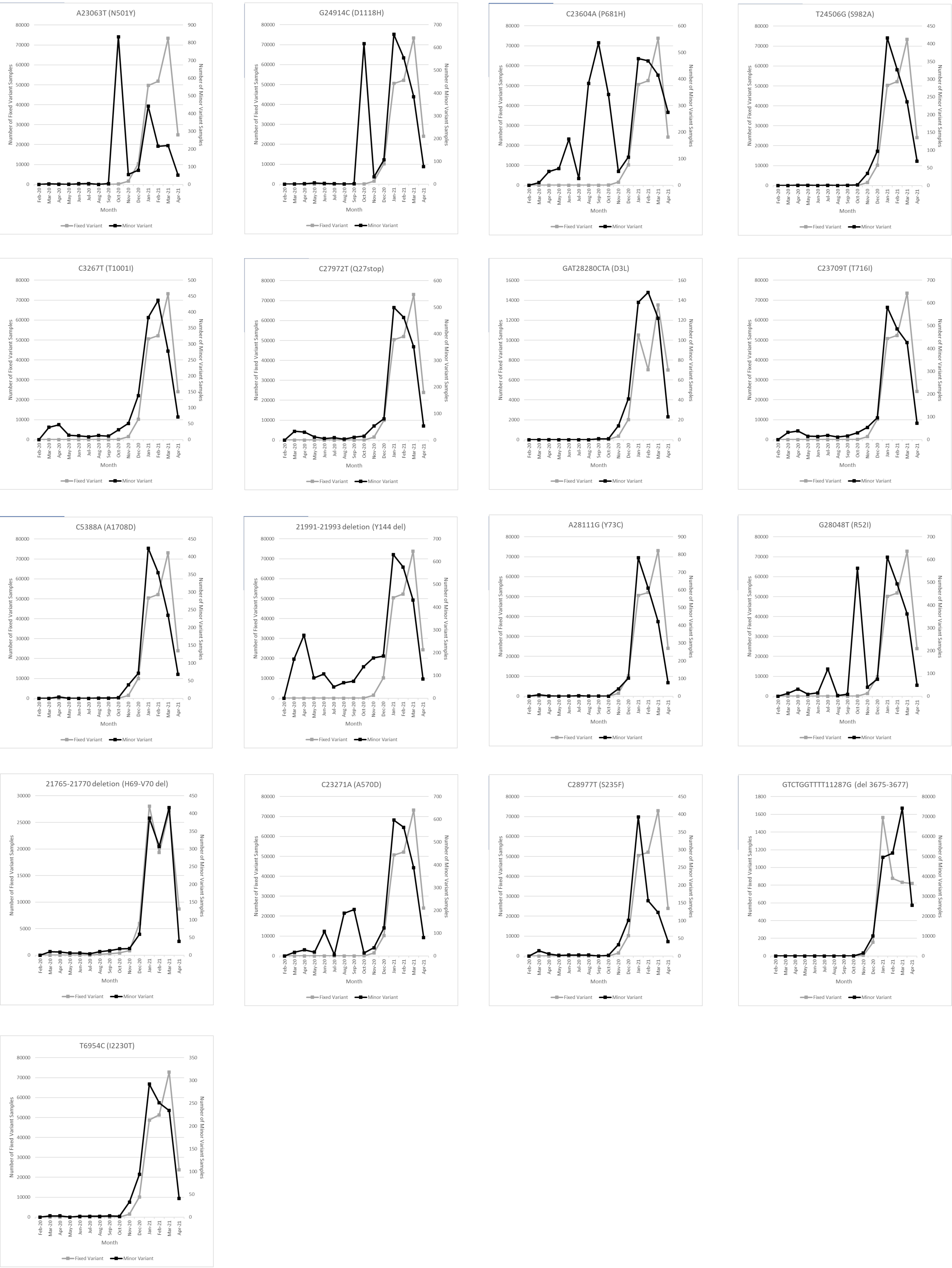

### Mutations Found in the Beta Variant (B.1.351 Lineage)

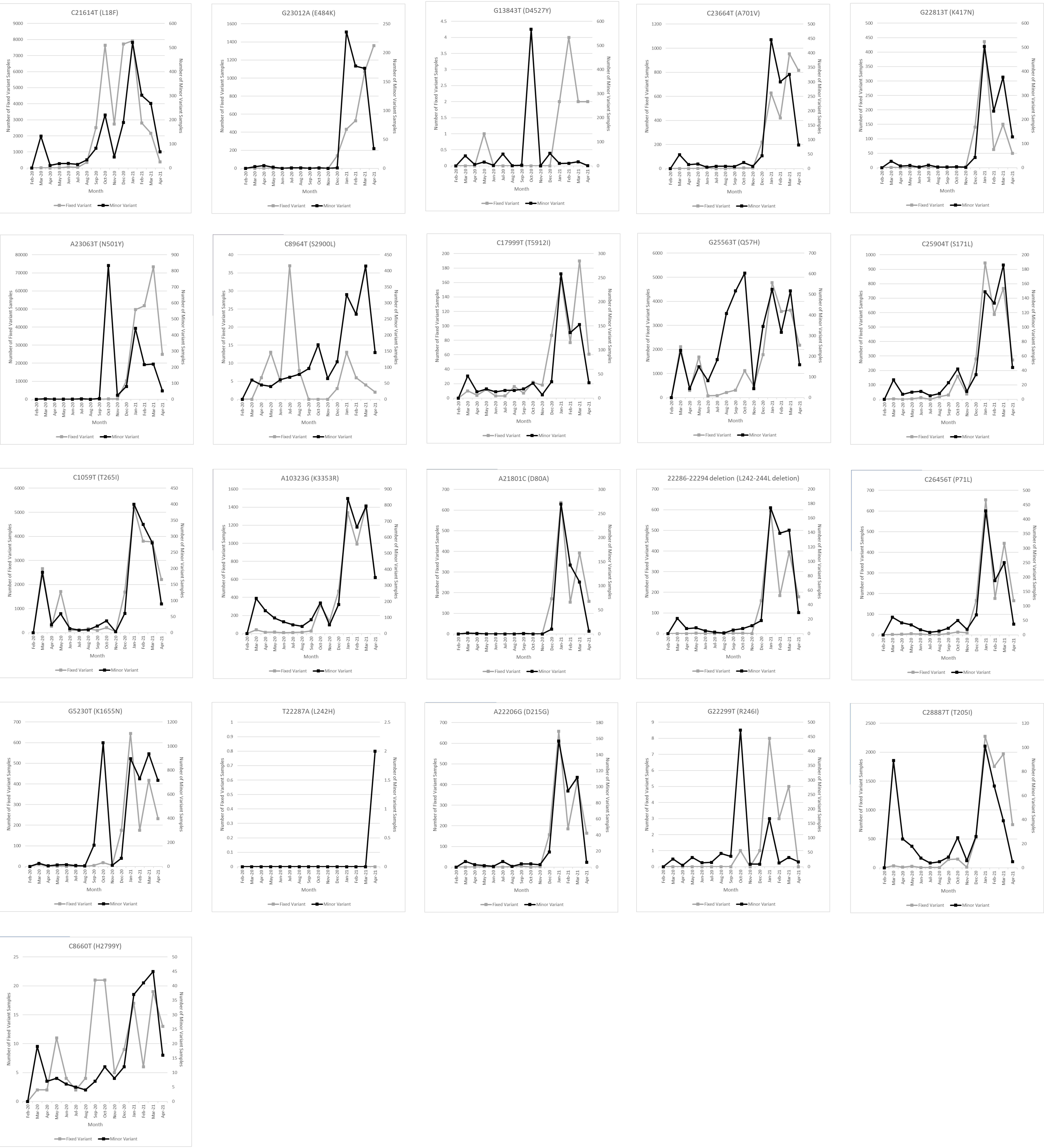

### Mutations Found in the Gamma Variant (P.1 Lineage)

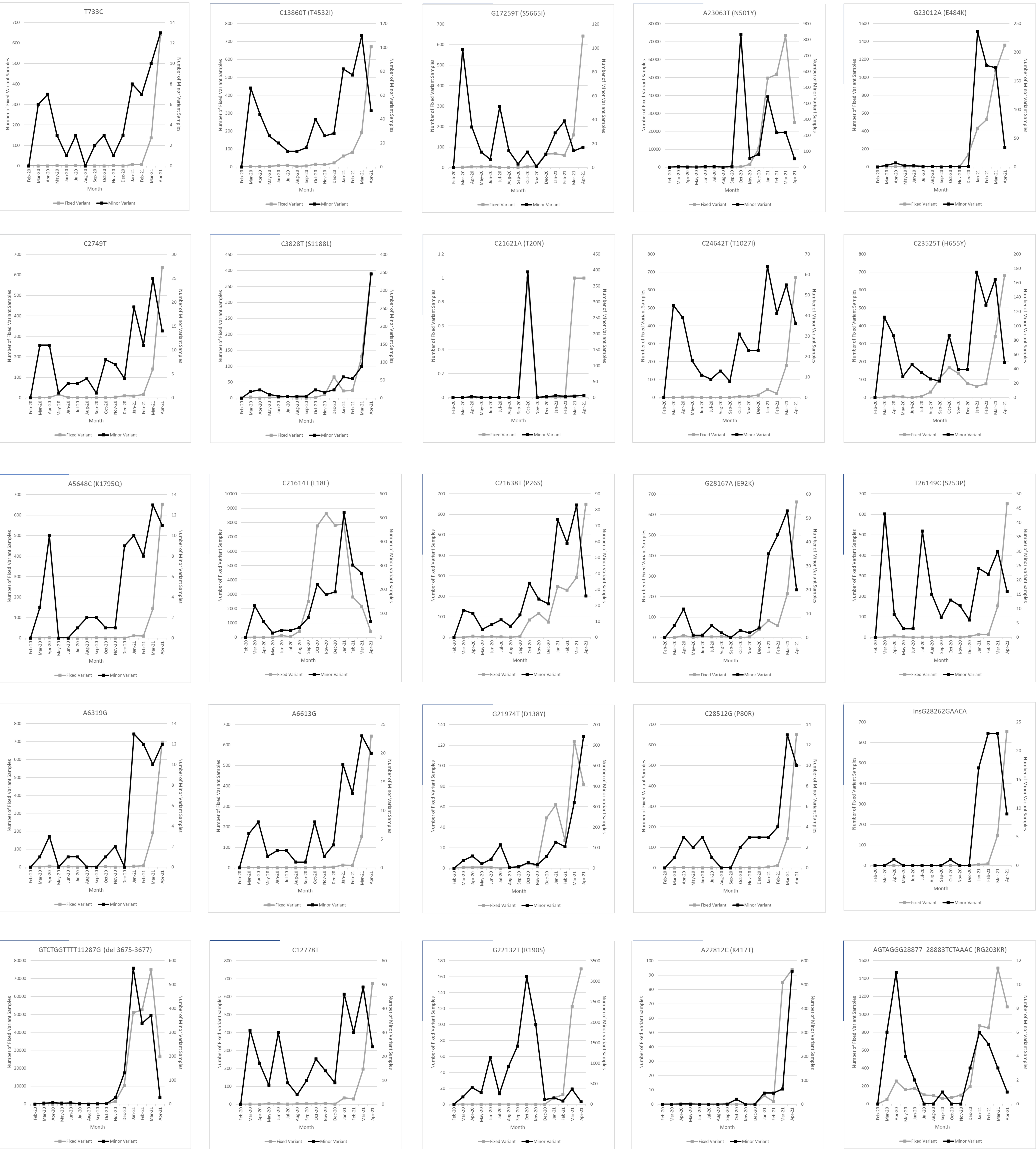

### Mutations Found in the Delta Variant (B.1.617.2 Lineage)

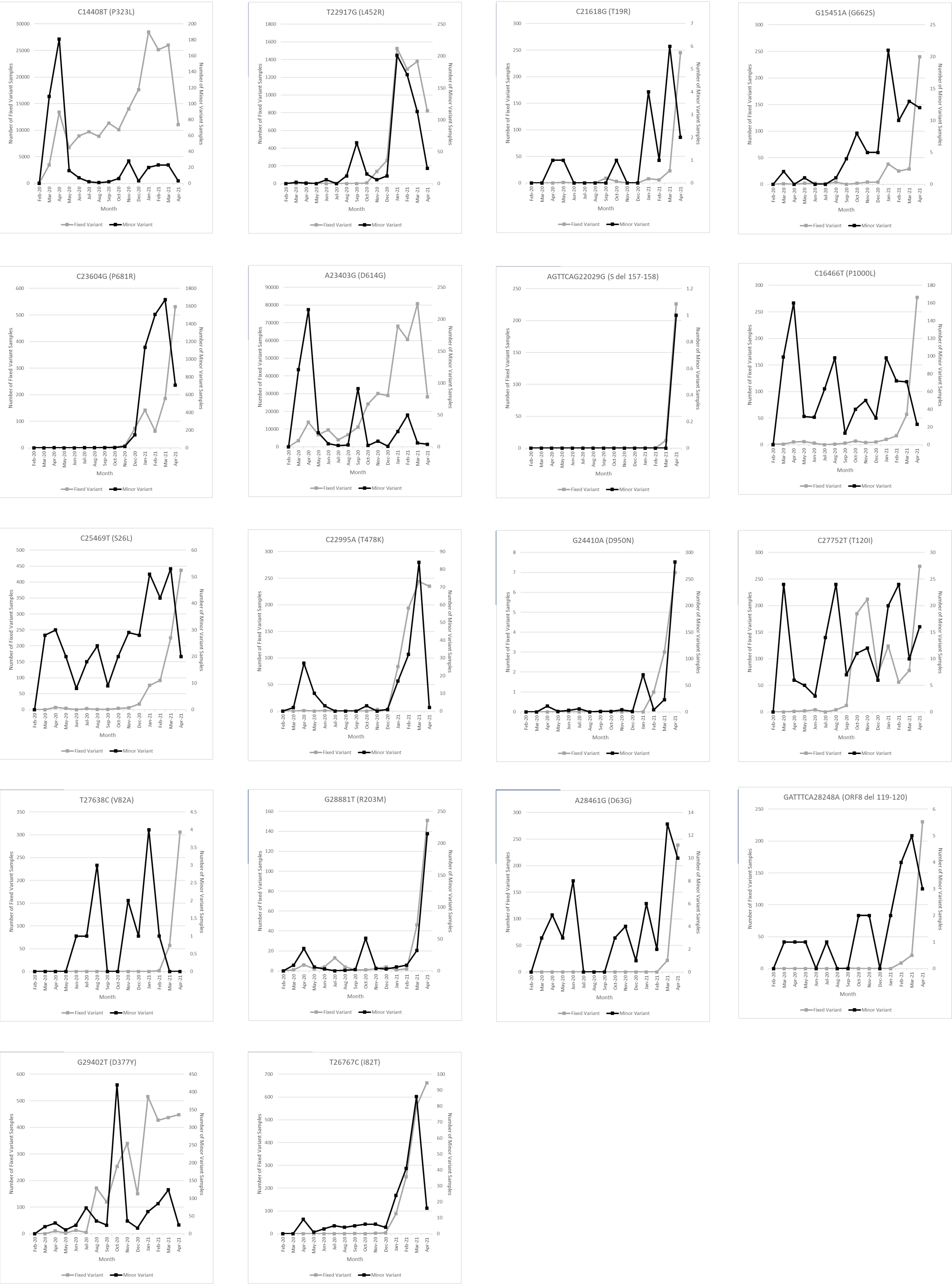

### Mutations Found in the Epsilon Variant (B.1.429 lineage)

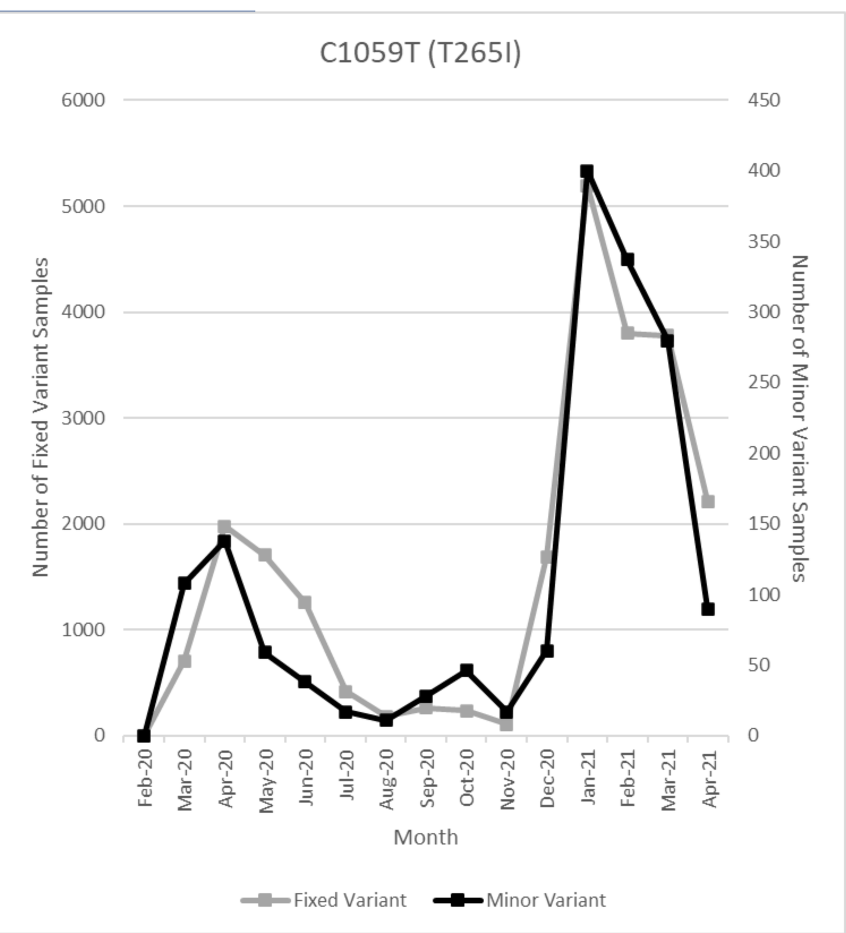


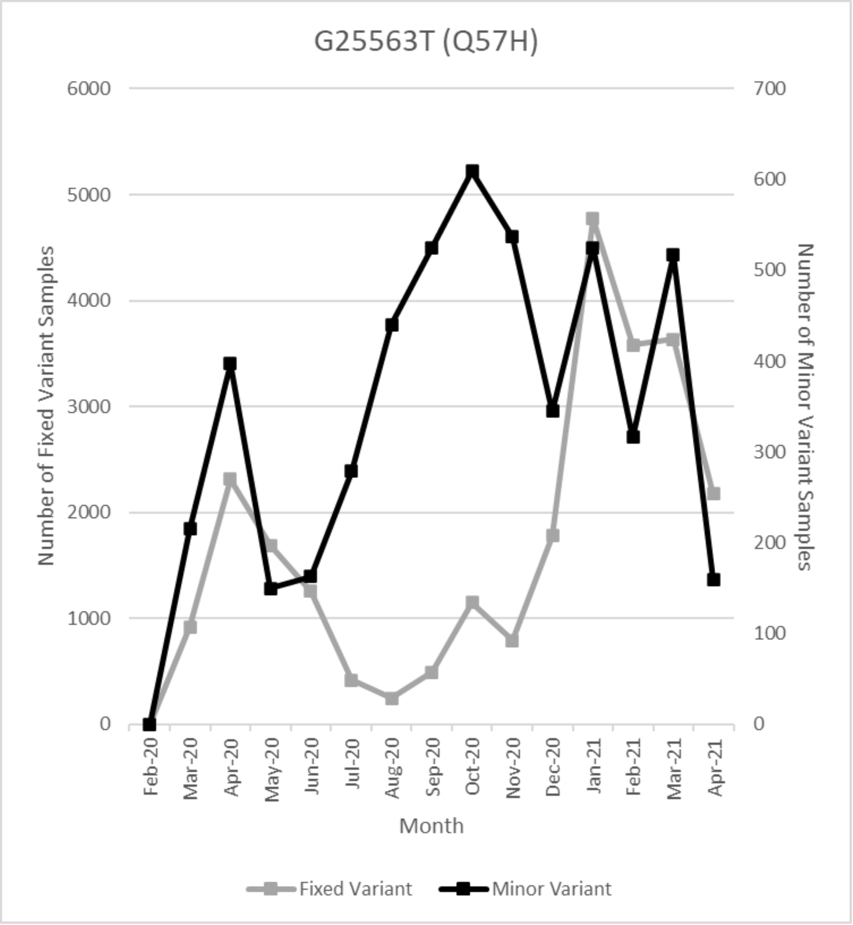


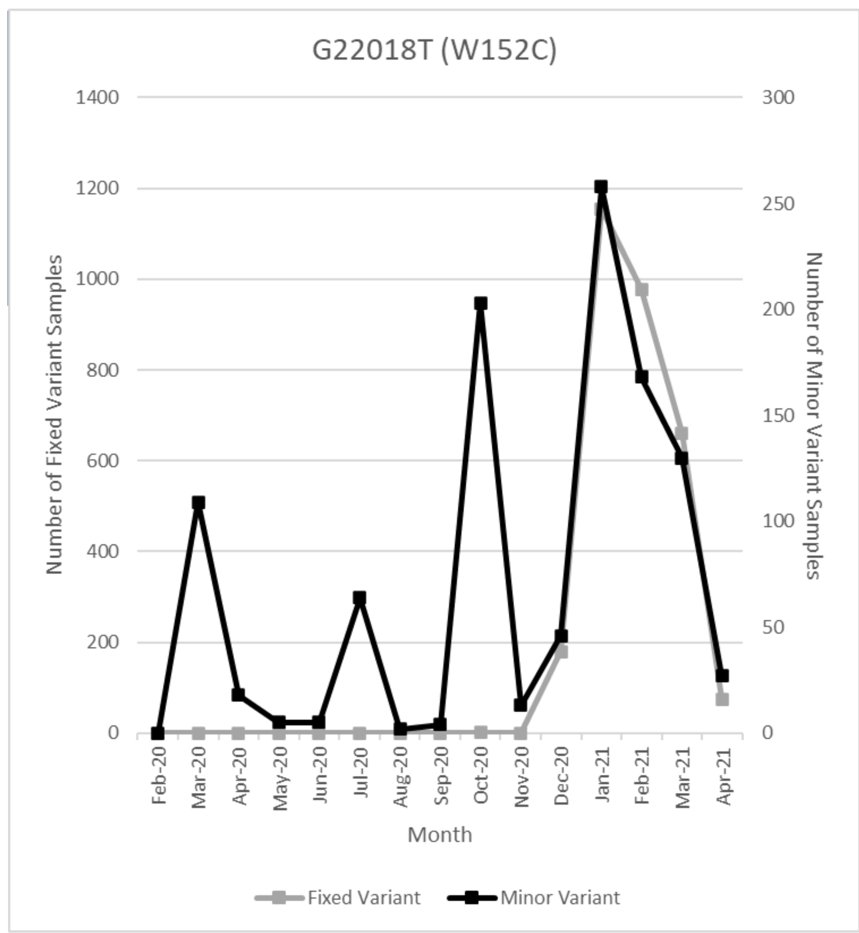


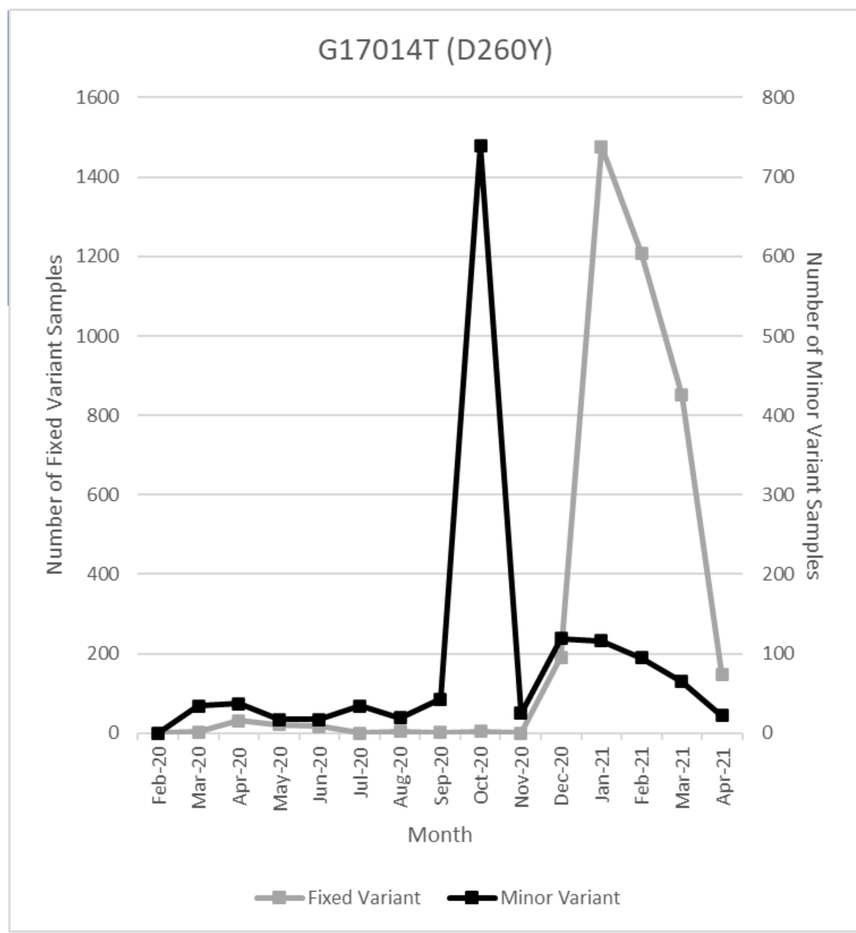


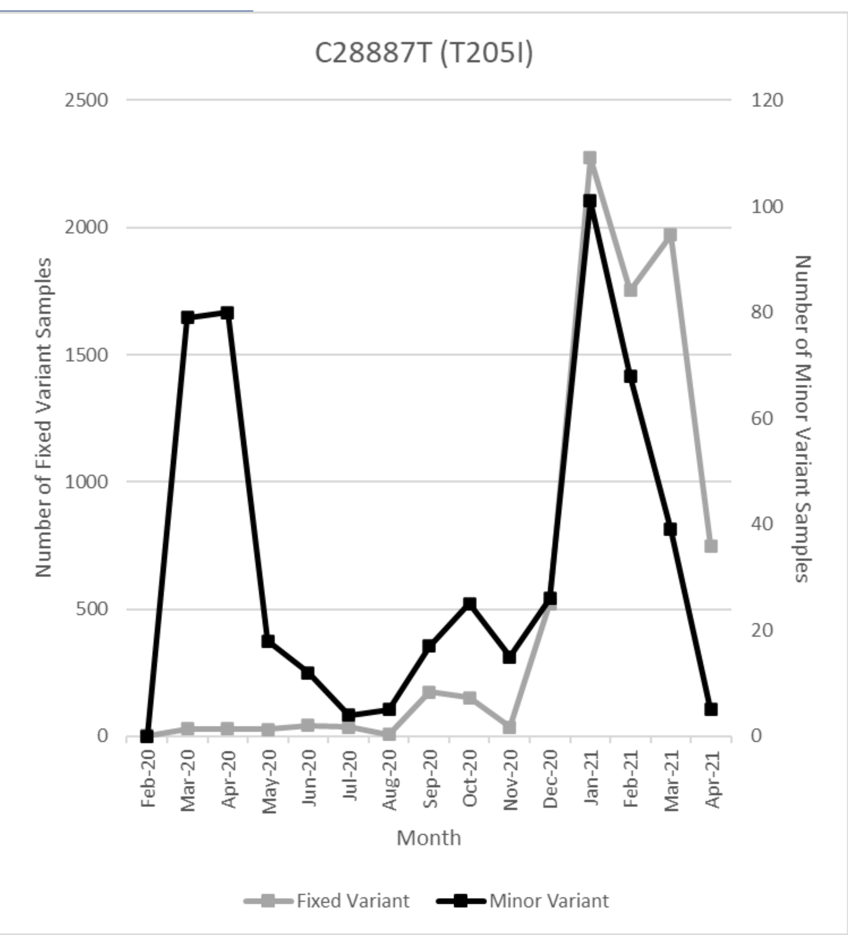


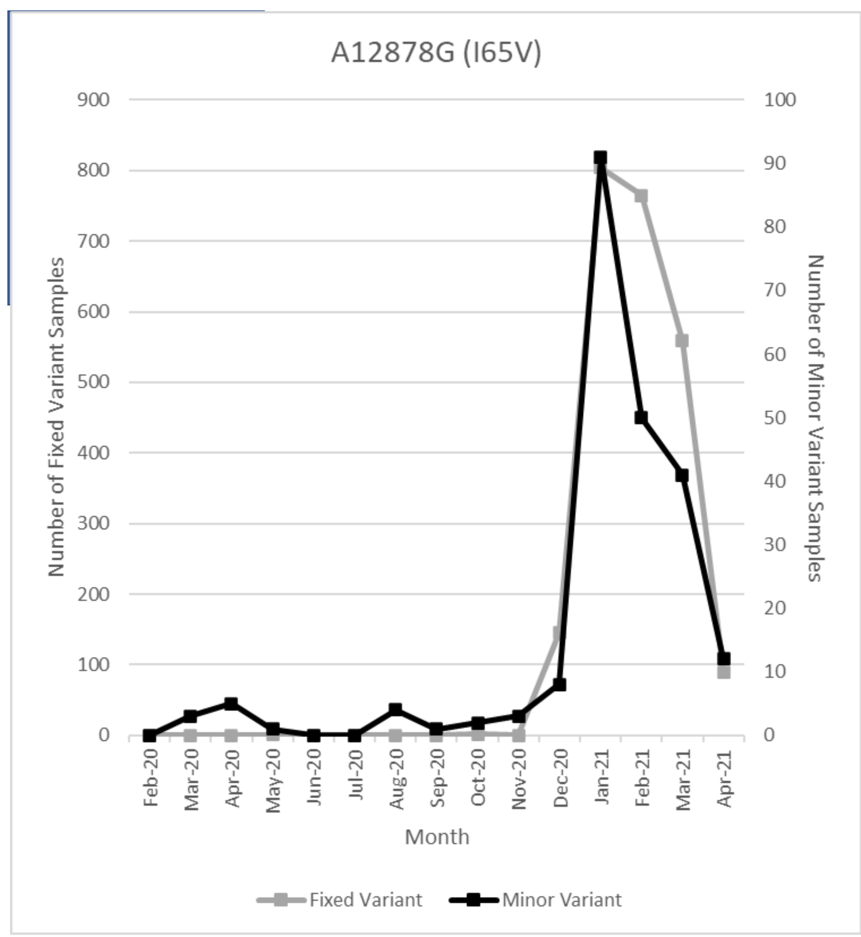


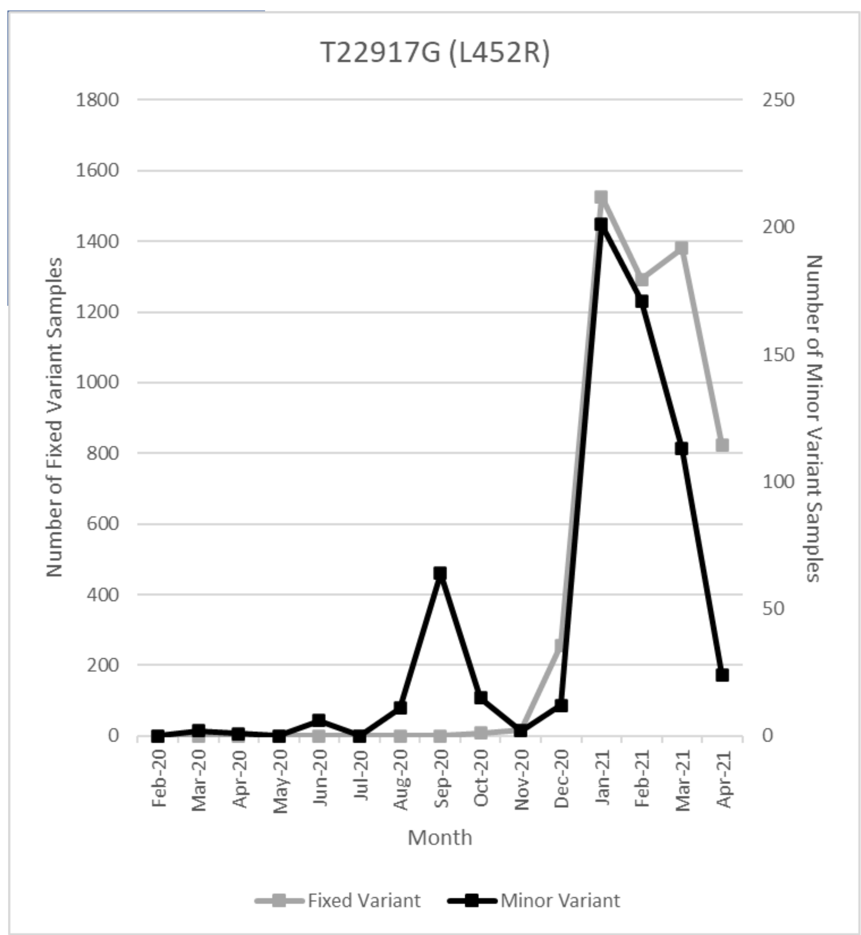


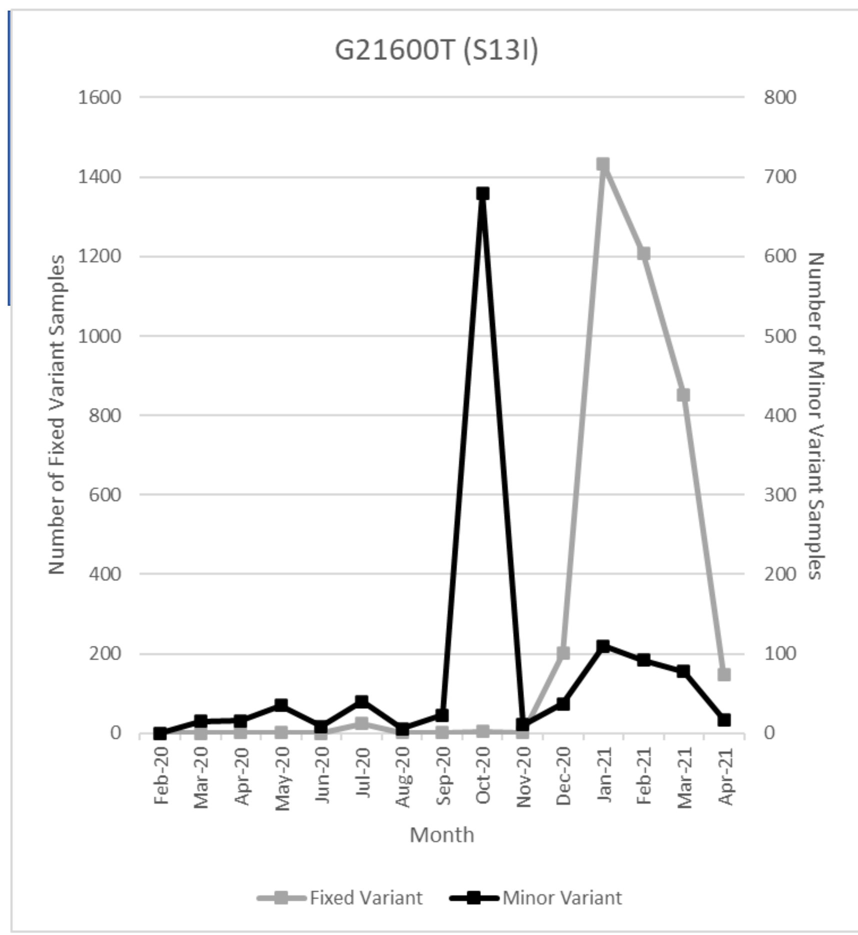


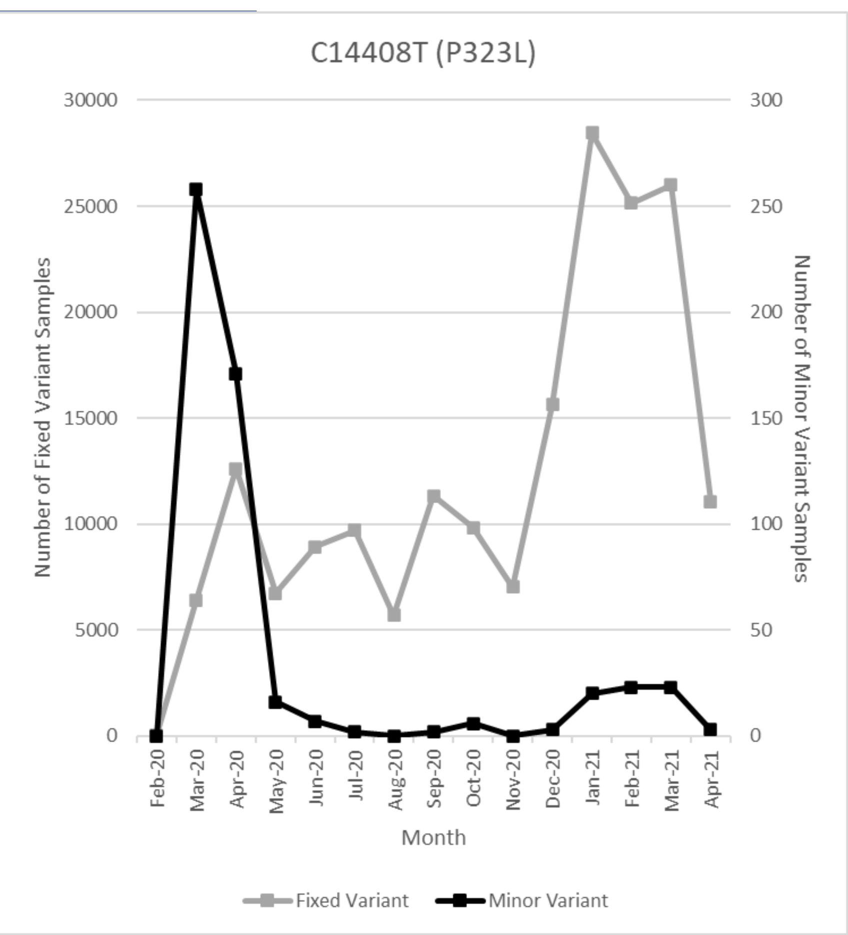


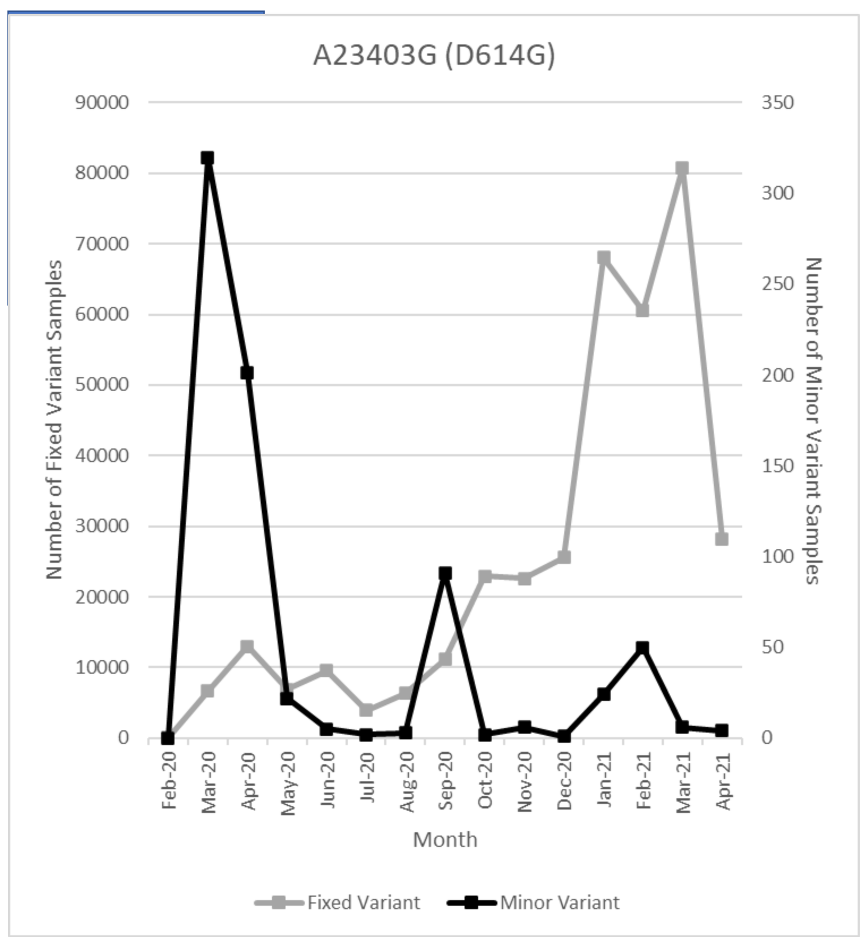

### Mutations Found in the Iota Variant (B.1.526 lineage)

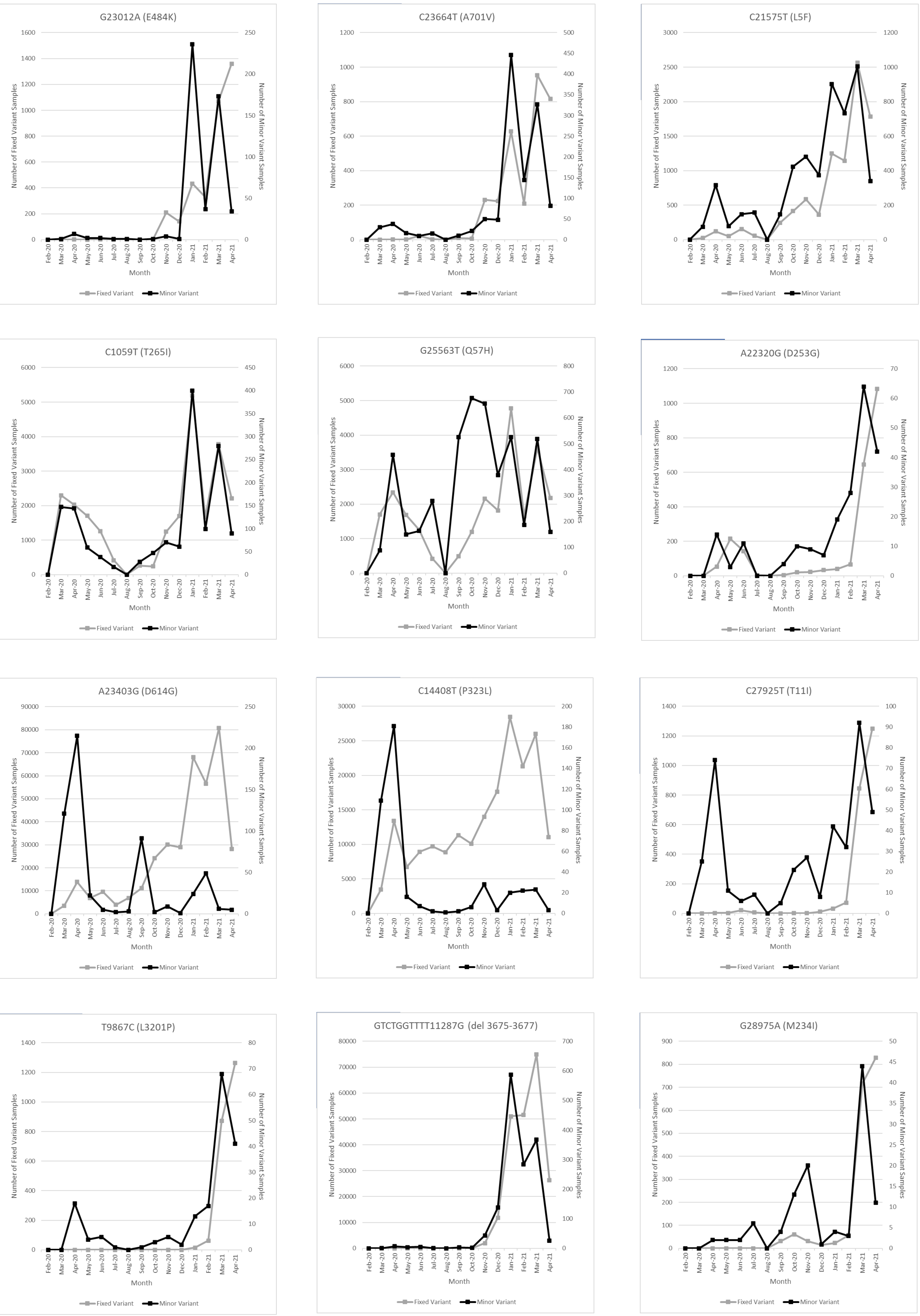

### Mutations Found in the Kappa Variant (B.1.617.1 Lineage)

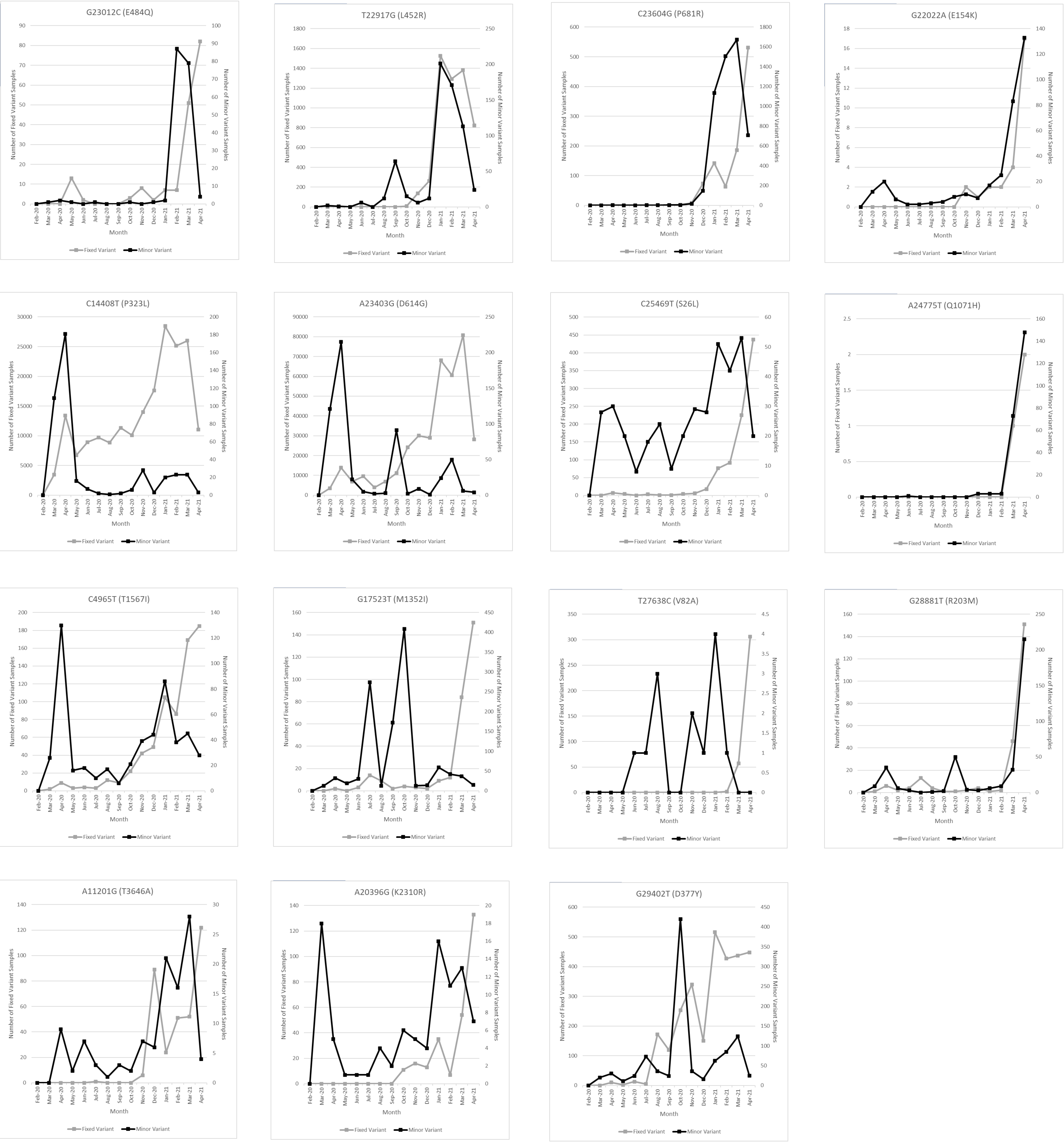
