## Supplementary figures and images for "Intrahost SARS-CoV-2 k-mer identification method (iSKIM) for rapid detection of mutations of concern reveals emergence of global mutation patterns"

### Figure S2

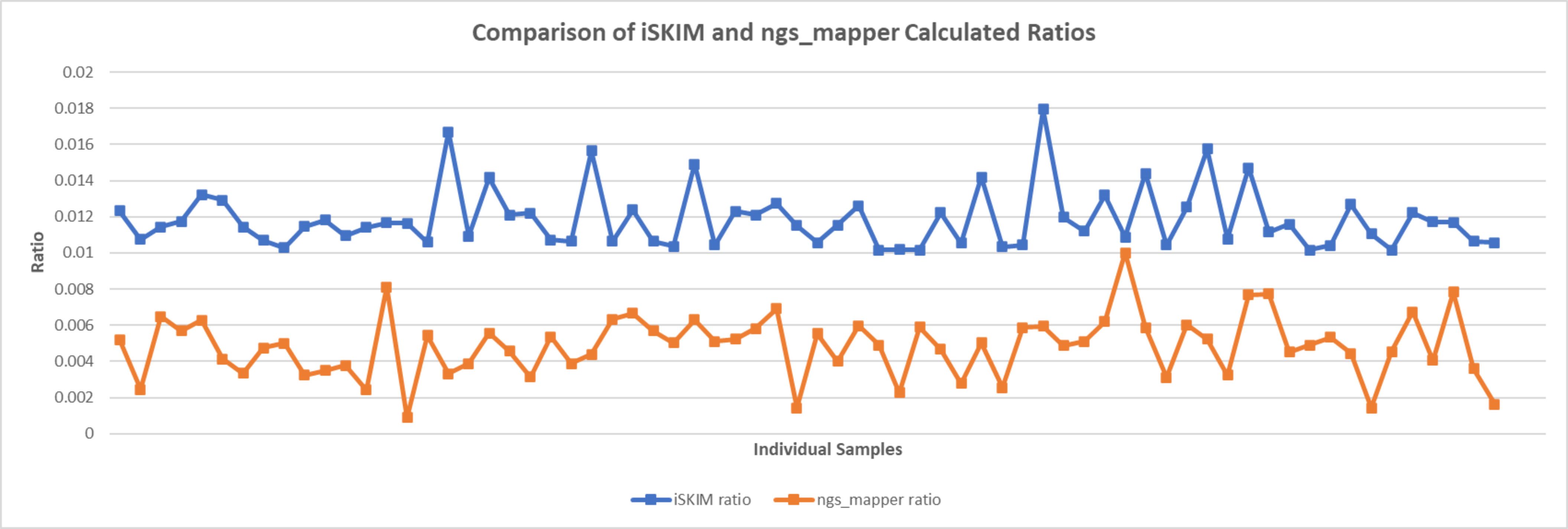

### Figure S3

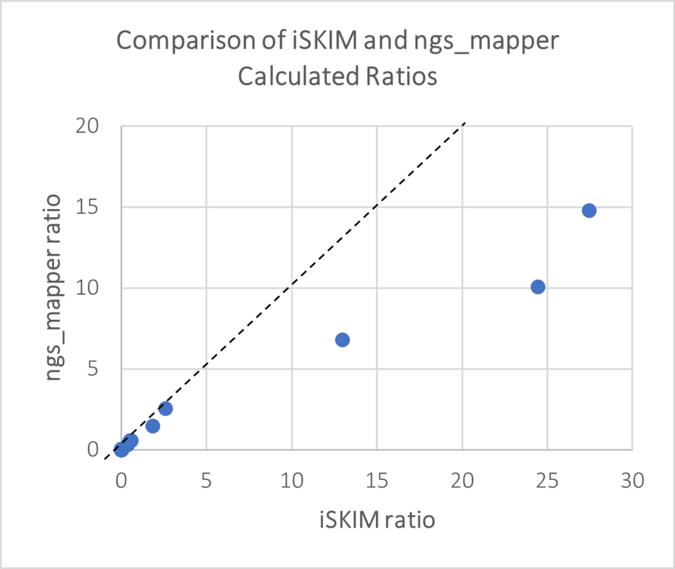

### Figure S4

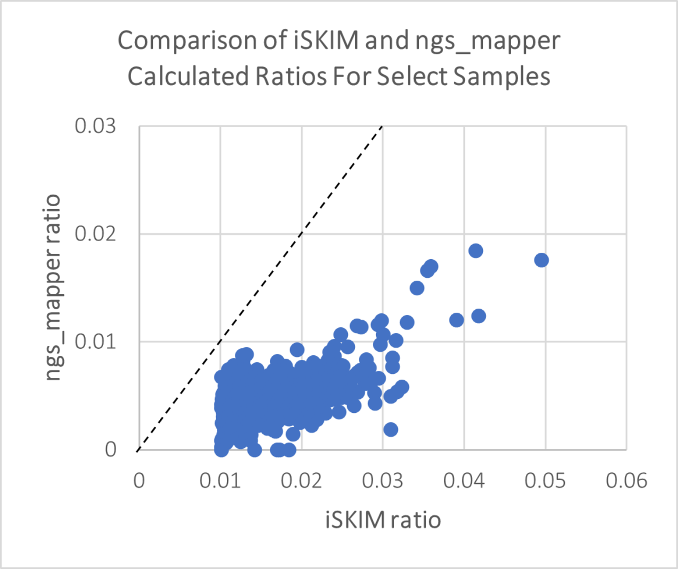

### Figure S5

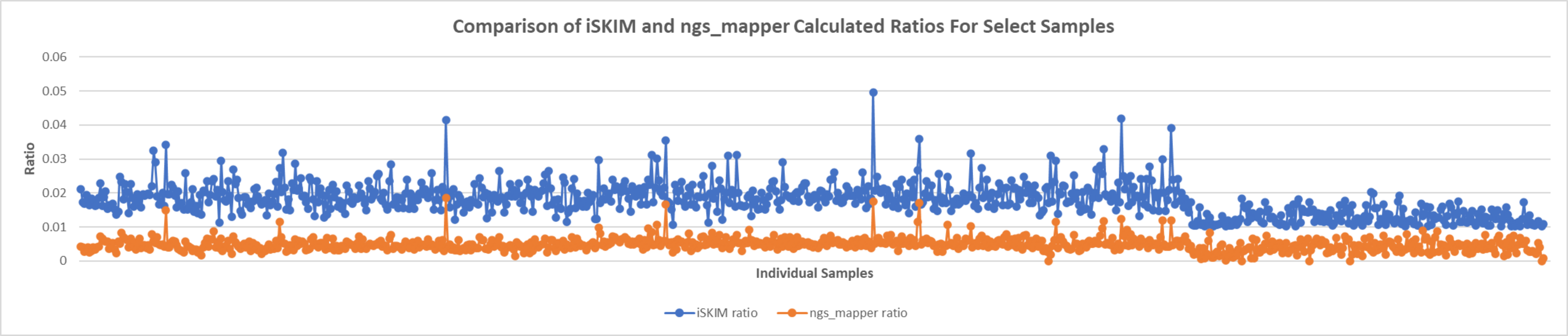

### Figure S6

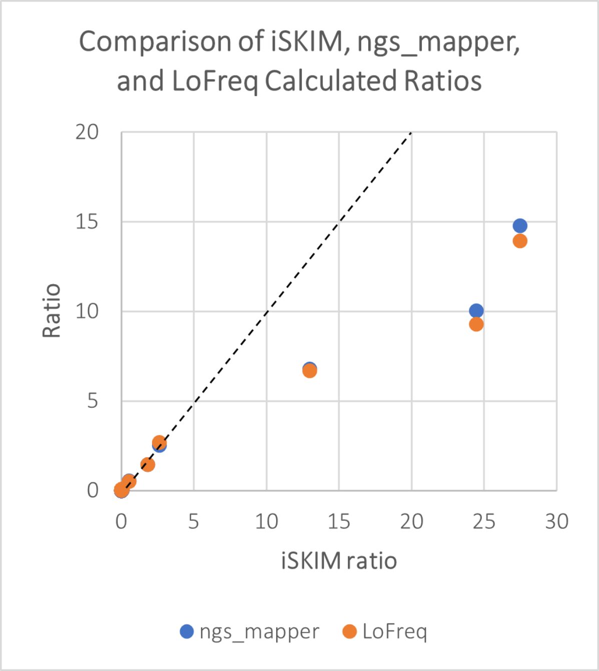

### Figure S7

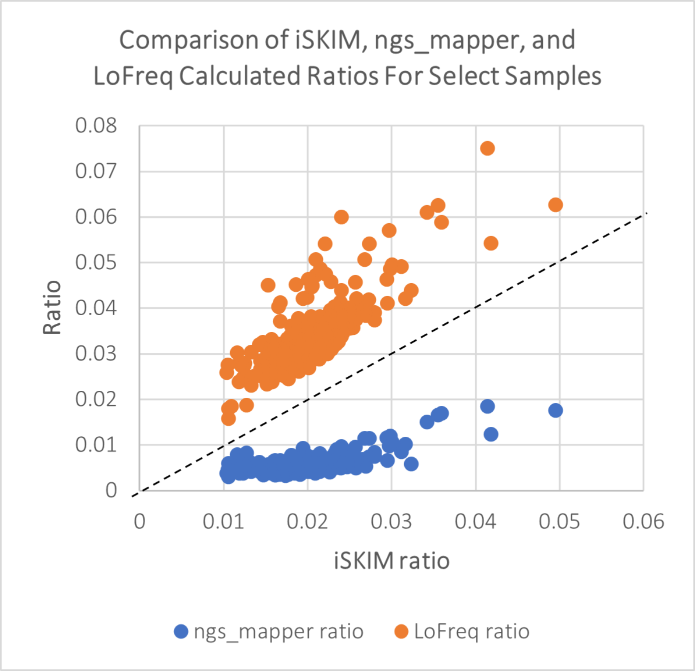

### Figure S8

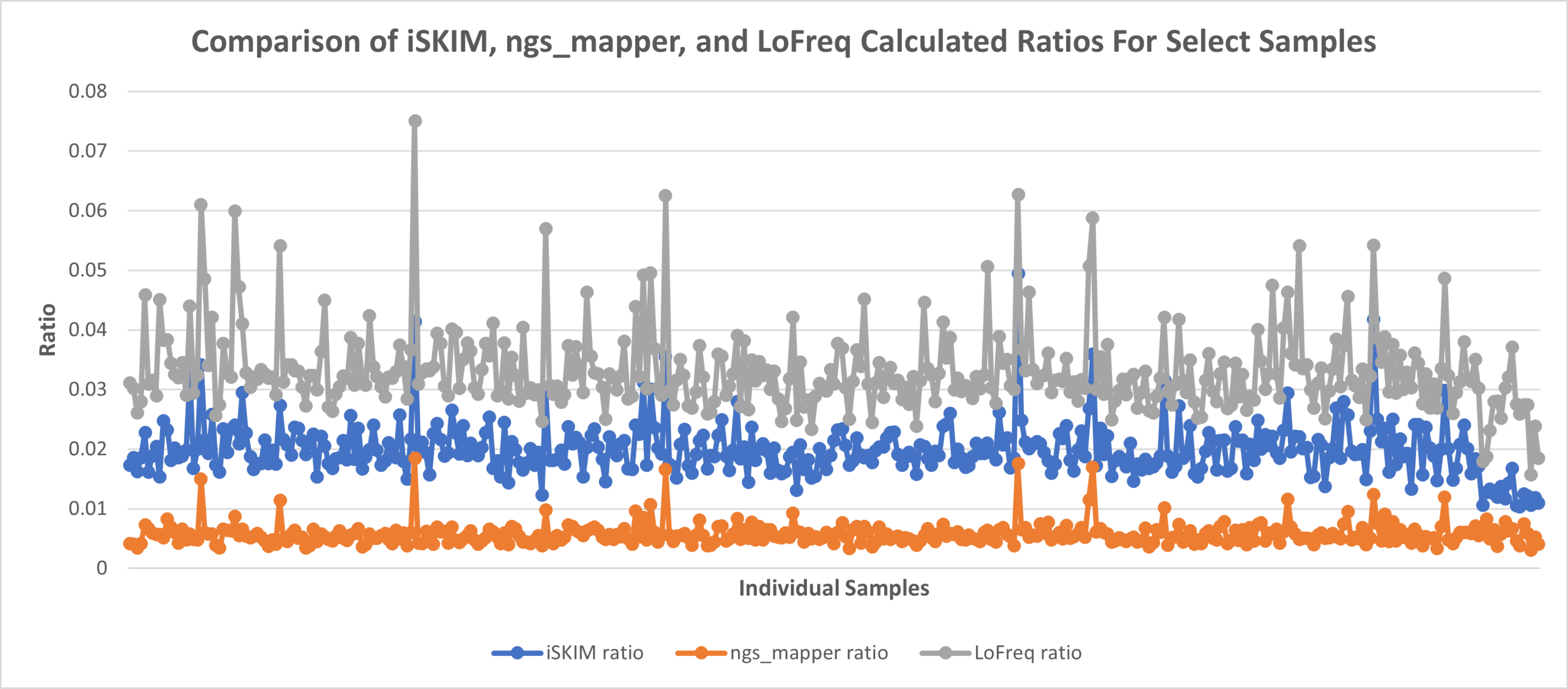
