## Supplementary material for "Intrahost SARS-CoV-2 k-mer identification method (iSKIM) for rapid detection of mutations of concern reveals emergence of global mutation patterns": Table S4

We gratefully acknowledge the following Authors from the Originating laboratories responsible for obtaining the specimens, as well as the Submitting laboratories where the genome data were generated and shared via GISAID, on which this research is based.

All Submitters of data may be contacted directly via www.gisaid.org

Authors are sorted alphabetically.

Acknowledgement EPI_SET Identifier: EPI_SET_20220218wq

| Accession ID | Originating Laboratory | Submitting Laboratory | Authors |
|---|---|---|---|
| EPI_ISL_611114, EPI_ISL_611142, EPI_ISL_611281, EPI_ISL_625032 | Lighthouse Lab in Alderley Park | Wellcome Sanger Institute for the COVID-19 Genomics UK (COG-UK) consortium | Cordelia Langford; David K. Jackson; Dominic Kwiatkowski; Ewan Harrison; Ian Johnston; Jacquelyn Wynn; John Sillitoe on behalf of the Wellcome Sanger Institute COVID-19 Surveillance Team (http://www.sanger.ac.uk/covid-team); Mairead Hyland; Roberto Amato; Sonia Goncalves; The Lighthouse Lab in Alderley Park and Alex Alderton |
| EPI_ISL_643565, EPI_ISL_672920, EPI_ISL_672966, EPI_ISL_673063, EPI_ISL_673100, EPI_ISL_673124, EPI_ISL_673133, EPI_ISL_673200 | | | |
| see above | Lighthouse Lab in Cambridge | Wellcome Sanger Institute for the COVID-19 Genomics UK (COG-UK) Consortium | Cordelia Langford; David K. Jackson; Dominic Kwiatkowski; Ewan Harrison; Ian Johnston; John Sillitoe on behalf of the Wellcome Sanger Institute COVID-19 Surveillance Team; Rob Howes; Roberto Amato; Sonia Goncalves; The Lighthouse Lab in Cambridge and Alex Alderton |
| EPI_ISL_608679 | Lighthouse Lab in Cambridge | Wellcome Sanger Institute for the COVID-19 Genomics UK (COG-UK) consortium | Cordelia Langford; David K. Jackson; Dominic Kwiatkowski; Ewan Harrison; Ian Johnston; John Sillitoe on behalf of the Wellcome Sanger Institute COVID-19 Surveillance Team; Rob Howes; Roberto Amato; Sonia Goncalves; The Lighthouse Lab in Cambridge and Alex Alderton |
| EPI_ISL_649688, EPI_ISL_649731, EPI_ISL_649739, EPI_ISL_662675, EPI_ISL_662677, EPI_ISL_662688, EPI_ISL_662704, EPI_ISL_662747, EPI_ISL_662754, EPI_ISL_662764, EPI_ISL_662769, EPI_ISL_662771, EPI_ISL_662789, EPI_ISL_662803, EPI_ISL_662823, EPI_ISL_662824, EPI_ISL_662834, EPI_ISL_662840, EPI_ISL_662849, EPI_ISL_662850, EPI_ISL_662875, EPI_ISL_662890, EPI_ISL_662897, EPI_ISL_662900, EPI_ISL_662901, EPI_ISL_662902, EPI_ISL_662905, EPI_ISL_662906, EPI_ISL_662915, EPI_ISL_662916, EPI_ISL_662917, EPI_ISL_662918, EPI_ISL_662920, EPI_ISL_662922, EPI_ISL_662923, EPI_ISL_662926, EPI_ISL_662927, EPI_ISL_662929, EPI_ISL_662931, EPI_ISL_662932, EPI_ISL_662934, EPI_ISL_662936, EPI_ISL_662938, EPI_ISL_662940, EPI_ISL_662943, EPI_ISL_662944, EPI_ISL_662946, EPI_ISL_662947, EPI_ISL_662950, EPI_ISL_662951, EPI_ISL_662954, EPI_ISL_662963, EPI_ISL_662966, EPI_ISL_662968, EPI_ISL_662975, EPI_ISL_662976, EPI_ISL_662984, EPI_ISL_662985, EPI_ISL_662986, EPI_ISL_662990, EPI_ISL_662992, EPI_ISL_662993, EPI_ISL_662994, EPI_ISL_662996, EPI_ISL_663000, EPI_ISL_663008, EPI_ISL_663010, EPI_ISL_663011, EPI_ISL_663012, EPI_ISL_663013, EPI_ISL_663016, EPI_ISL_663018, EPI_ISL_663020, EPI_ISL_663021, EPI_ISL_663023, EPI_ISL_663024, EPI_ISL_663025, EPI_ISL_663027, EPI_ISL_663031, EPI_ISL_663032, EPI_ISL_663036, EPI_ISL_663037, EPI_ISL_663038, EPI_ISL_663040, EPI_ISL_663042, EPI_ISL_663047, EPI_ISL_663049, EPI_ISL_663050, EPI_ISL_663051, EPI_ISL_663053, EPI_ISL_663054, EPI_ISL_663058, EPI_ISL_663060, EPI_ISL_663061, EPI_ISL_663065, EPI_ISL_663066, EPI_ISL_663070, EPI_ISL_663071, EPI_ISL_663075, EPI_ISL_663077, EPI_ISL_663078, EPI_ISL_663080, EPI_ISL_663081, EPI_ISL_663082, EPI_ISL_663089, EPI_ISL_663093, EPI_ISL_663099, EPI_ISL_663101, EPI_ISL_663103, EPI_ISL_663105, EPI_ISL_663107, EPI_ISL_663109, EPI_ISL_663110, EPI_ISL_663113, EPI_ISL_663115, EPI_ISL_663118, EPI_ISL_663121, EPI_ISL_663124, EPI_ISL_663129, EPI_ISL_663130, EPI_ISL_663131, EPI_ISL_663133, EPI_ISL_663135, EPI_ISL_663139, EPI_ISL_663140, EPI_ISL_663142, EPI_ISL_663145, EPI_ISL_663146, EPI_ISL_663147, EPI_ISL_663149, EPI_ISL_663151, EPI_ISL_663158, EPI_ISL_663160, EPI_ISL_663167, EPI_ISL_663168, EPI_ISL_663171, EPI_ISL_663172, EPI_ISL_663173, EPI_ISL_663174, EPI_ISL_663175, EPI_ISL_663177, EPI_ISL_663179, EPI_ISL_663181, EPI_ISL_663185, EPI_ISL_663189, EPI_ISL_663191, EPI_ISL_663192, EPI_ISL_663195, EPI_ISL_663201, EPI_ISL_663202, EPI_ISL_663203, EPI_ISL_663204, EPI_ISL_676446, EPI_ISL_676451, EPI_ISL_676461, EPI_ISL_676462, EPI_ISL_676464, EPI_ISL_676465, EPI_ISL_676467, EPI_ISL_676469, EPI_ISL_676470, EPI_ISL_676472, EPI_ISL_676473, EPI_ISL_676475, EPI_ISL_676477, EPI_ISL_676482, EPI_ISL_721562 | | | |
| see above | Lighthouse Lab in Glasgow | Wellcome Sanger Institute for the COVID-19 Genomics UK (COG-UK) Consortium | Anna Dominiczak and Alex Alderton; Carol Clugston; Cordelia Langford; David Gray; David K. Jackson; Dominic Kwiatkowski; Ewan Harrison; Harper VanSteenhouse; Ian Johnston; John Sillitoe on behalf of the Wellcome Sanger Institute COVID-19 Surveillance Team; John Sillitoe on behalf of the Wellcome Sanger Institute COVID-19 Surveillance Team (http://www.sanger.ac.uk/covid-team); Roberto Amato; Sonia Goncalves; Yumi Kasai |
| EPI_ISL_611477, EPI_ISL_625328, EPI_ISL_633477, EPI_ISL_634070, EPI_ISL_634072, EPI_ISL_634075, EPI_ISL_634097, EPI_ISL_634114, EPI_ISL_634117, EPI_ISL_634135, EPI_ISL_634207, EPI_ISL_634224, EPI_ISL_634234, EPI_ISL_634235, EPI_ISL_634240 | | | |
| see above | Lighthouse Lab in Glasgow | Wellcome Sanger Institute for the COVID-19 Genomics UK (COG-UK) consortium | Anna Dominiczak and Alex Alderton; Carol Clugston; Cordelia Langford; David Gray; David K. Jackson; Dominic Kwiatkowski; Ewan Harrison; Harper VanSteenhouse; Ian Johnston; John Sillitoe on behalf of the Wellcome Sanger Institute COVID-19 Surveillance Team; John Sillitoe on behalf of the Wellcome Sanger Institute COVID-19 Surveillance Team (http://www.sanger.ac.uk/covid-team); Roberto Amato; Sonia Goncalves; Yumi Kasai |
| EPI_ISL_598249, EPI_ISL_598305, EPI_ISL_598330, EPI_ISL_598331, EPI_ISL_598408, EPI_ISL_598433, EPI_ISL_598606, EPI_ISL_598638, EPI_ISL_598667, EPI_ISL_598821, EPI_ISL_609230, EPI_ISL_609267, EPI_ISL_609302 | | | |
| see above | Lighthouse Lab in Milton Keynes | Wellcome Sanger Institute for the COVID-19 Genomics UK (COG-UK) Consortium | Cordelia Langford; David K. Jackson; Dominic Kwiatkowski; Ewan Harrison; Ian Johnston; John Sillitoe on behalf of the Wellcome Sanger Institute COVID-19 Surveillance Team; John Sillitoe on behalf of the Wellcome Sanger Institute COVID-19 Surveillance Team (http://www.sanger.ac.uk/covid-team); Roberto Amato; Sonia Goncalves; The Lighthouse Lab in Milton Keynes and Alex Alderton |
| EPI_ISL_598250, EPI_ISL_598251, EPI_ISL_598252, EPI_ISL_598253, EPI_ISL_598254, EPI_ISL_598256, EPI_ISL_598257, EPI_ISL_598258, EPI_ISL_598259, EPI_ISL_598260, EPI_ISL_598261, EPI_ISL_598262, EPI_ISL_598263, EPI_ISL_598264, EPI_ISL_598265, EPI_ISL_598266, EPI_ISL_598267, EPI_ISL_598268, EPI_ISL_598270, EPI_ISL_598271, EPI_ISL_598272, EPI_ISL_598273, EPI_ISL_598274, EPI_ISL_598275, EPI_ISL_598276, EPI_ISL_598277, EPI_ISL_598278, EPI_ISL_598279, EPI_ISL_598280, EPI_ISL_598281, EPI_ISL_598282, EPI_ISL_598283, EPI_ISL_598284, EPI_ISL_598286, EPI_ISL_598287, EPI_ISL_598288, EPI_ISL_598289, EPI_ISL_598290, EPI_ISL_598291, EPI_ISL_598292, EPI_ISL_598293, EPI_ISL_598294, EPI_ISL_598295, EPI_ISL_598296, EPI_ISL_598297, EPI_ISL_598298, EPI_ISL_598299, EPI_ISL_598300, EPI_ISL_598301, EPI_ISL_598302, EPI_ISL_598303, EPI_ISL_598304, EPI_ISL_598306, EPI_ISL_598307, EPI_ISL_598308, EPI_ISL_598309, EPI_ISL_598310, EPI_ISL_598311, EPI_ISL_598313, EPI_ISL_598314, EPI_ISL_598315, EPI_ISL_598316, EPI_ISL_598317, EPI_ISL_598318, EPI_ISL_598319, EPI_ISL_598320, EPI_ISL_598321, EPI_ISL_598322, EPI_ISL_598323, EPI_ISL_598324, EPI_ISL_598325, EPI_ISL_598326, EPI_ISL_598327, EPI_ISL_598328, EPI_ISL_598329, EPI_ISL_598332, EPI_ISL_598333, EPI_ISL_598334, EPI_ISL_598335, EPI_ISL_598336, EPI_ISL_598337, EPI_ISL_598338, EPI_ISL_598339, EPI_ISL_598340, EPI_ISL_598341, EPI_ISL_598342, EPI_ISL_598343, EPI_ISL_598344, EPI_ISL_598345, EPI_ISL_598346, EPI_ISL_598347, EPI_ISL_598348, EPI_ISL_598349, EPI_ISL_598350, EPI_ISL_598352, EPI_ISL_598353, EPI_ISL_598354, EPI_ISL_598355, EPI_ISL_598356, EPI_ISL_598357, EPI_ISL_598358, EPI_ISL_598359, EPI_ISL_598360, EPI_ISL_598362, EPI_ISL_598363, EPI_ISL_598365, EPI_ISL_598366, EPI_ISL_598367, EPI_ISL_598369, EPI_ISL_598370, EPI_ISL_598371, EPI_ISL_598372, EPI_ISL_598373, EPI_ISL_598374, EPI_ISL_598375, EPI_ISL_598376, EPI_ISL_598377, EPI_ISL_598378, EPI_ISL_598379, EPI_ISL_598380, EPI_ISL_598381, EPI_ISL_598382, EPI_ISL_598383, EPI_ISL_598385, EPI_ISL_598386, EPI_ISL_598387, EPI_ISL_598389, EPI_ISL_598390, EPI_ISL_598391, EPI_ISL_598392, EPI_ISL_598393, EPI_ISL_598394, EPI_ISL_598395, EPI_ISL_598396, EPI_ISL_598397, EPI_ISL_598398, EPI_ISL_598399, EPI_ISL_598400, EPI_ISL_598401, EPI_ISL_598402, EPI_ISL_598403, EPI_ISL_598404, EPI_ISL_598405, EPI_ISL_598406, EPI_ISL_598407, EPI_ISL_598409, EPI_ISL_598410, EPI_ISL_598411, EPI_ISL_598412, EPI_ISL_598413, EPI_ISL_598414, EPI_ISL_598415, EPI_ISL_598416, EPI_ISL_598417, EPI_ISL_598418, EPI_ISL_598419, EPI_ISL_598420, EPI_ISL_598421, EPI_ISL_598422, EPI_ISL_598423, EPI_ISL_598424, EPI_ISL_598425, EPI_ISL_598426, EPI_ISL_598427, EPI_ISL_598428, EPI_ISL_598429, EPI_ISL_598430, EPI_ISL_598431, EPI_ISL_598432, EPI_ISL_598434, EPI_ISL_598435, EPI_ISL_598437, EPI_ISL_598438, EPI_ISL_598439, EPI_ISL_598440, EPI_ISL_598441, EPI_ISL_598442, EPI_ISL_598443, EPI_ISL_598444, EPI_ISL_598445, EPI_ISL_598446, EPI_ISL_598447, EPI_ISL_598448, EPI_ISL_598449, EPI_ISL_598450, EPI_ISL_598451, EPI_ISL_598452, EPI_ISL_598453, EPI_ISL_598454, EPI_ISL_598455, EPI_ISL_598456, EPI_ISL_598457, EPI_ISL_598458, EPI_ISL_598459, EPI_ISL_598460, EPI_ISL_598461, EPI_ISL_598462, EPI_ISL_598463, EPI_ISL_598464, EPI_ISL_598465, EPI_ISL_598466, EPI_ISL_598467, EPI_ISL_598468, EPI_ISL_598469, EPI_ISL_598470, EPI_ISL_598471, EPI_ISL_598472, EPI_ISL_598473, EPI_ISL_598474, EPI_ISL_598475, EPI_ISL_598476, EPI_ISL_598477, EPI_ISL_598478, EPI_ISL_598479, EPI_ISL_598480, EPI_ISL_598481, EPI_ISL_598482, EPI_ISL_598483, EPI_ISL_598484, EPI_ISL_598485, EPI_ISL_598486, EPI_ISL_598487, EPI_ISL_598488, EPI_ISL_598489, EPI_ISL_598490, EPI_ISL_598491, EPI_ISL_598492, EPI_ISL_598493, EPI_ISL_598494, EPI_ISL_598495, EPI_ISL_598496, EPI_ISL_598497, EPI_ISL_598498, EPI_ISL_598499, EPI_ISL_598500, EPI_ISL_598501, EPI_ISL_598502, EPI_ISL_598503, EPI_ISL_598505, EPI_ISL_598506, EPI_ISL_598507, EPI_ISL_598508, EPI_ISL_598509, EPI_ISL_598510, EPI_ISL_598511, EPI_ISL_598512, EPI_ISL_598513, EPI_ISL_598515, EPI_ISL_598516, EPI_ISL_598517, EPI_ISL_598518, EPI_ISL_598519, EPI_ISL_598520, EPI_ISL_598521, EPI_ISL_598523, EPI_ISL_598524, EPI_ISL_598525, EPI_ISL_598526, EPI_ISL_598527, EPI_ISL_598528, EPI_ISL_598529, EPI_ISL_598530, EPI_ISL_598531, EPI_ISL_598532, EPI_ISL_598533, EPI_ISL_598534, EPI_ISL_598536, EPI_ISL_598537, EPI_ISL_598538, EPI_ISL_598539, EPI_ISL_598540, EPI_ISL_598541, EPI_ISL_598542, EPI_ISL_598543, EPI_ISL_598544, EPI_ISL_598546, EPI_ISL_598547, EPI_ISL_598548, EPI_ISL_598549, EPI_ISL_598550, EPI_ISL_598551, EPI_ISL_598552, EPI_ISL_598553, EPI_ISL_598554, EPI_ISL_598555, EPI_ISL_598556, EPI_ISL_598557, EPI_ISL_598558, EPI_ISL_598559, EPI_ISL_598560, EPI_ISL_598562, EPI_ISL_598563, EPI_ISL_598565, EPI_ISL_598566, EPI_ISL_598567, EPI_ISL_598569, EPI_ISL_598570, EPI_ISL_598571, EPI_ISL_598573, EPI_ISL_598575, EPI_ISL_598576, EPI_ISL_598577, EPI_ISL_598578, EPI_ISL_598579, EPI_ISL_598580, EPI_ISL_598581, EPI_ISL_598582, EPI_ISL_598583, EPI_ISL_598584, EPI_ISL_598585, EPI_ISL_598586, EPI_ISL_598587, EPI_ISL_598588, EPI_ISL_598589, EPI_ISL_598590, EPI_ISL_598591, EPI_ISL_598592, EPI_ISL_598593, EPI_ISL_598594, EPI_ISL_598595, EPI_ISL_598596, EPI_ISL_598597, EPI_ISL_598598, EPI_ISL_598599, EPI_ISL_598600, EPI_ISL_598601, EPI_ISL_598602, EPI_ISL_598604, EPI_ISL_598605, EPI_ISL_598607, EPI_ISL_598608, EPI_ISL_598609, EPI_ISL_598610, EPI_ISL_598611, EPI_ISL_598612, EPI_ISL_598613, EPI_ISL_598614, EPI_ISL_598615, EPI_ISL_598616, EPI_ISL_598617, EPI_ISL_598618, EPI_ISL_598619, EPI_ISL_598620, EPI_ISL_598621, EPI_ISL_598622, EPI_ISL_598623, EPI_ISL_598624, EPI_ISL_598625, EPI_ISL_598626, EPI_ISL_598627, EPI_ISL_598628, EPI_ISL_598629, EPI_ISL_598630, EPI_ISL_598631, EPI_ISL_598632, EPI_ISL_598633, EPI_ISL_598634, EPI_ISL_598636, EPI_ISL_598637, EPI_ISL_598639, EPI_ISL_598640, EPI_ISL_598641, EPI_ISL_598642, EPI_ISL_598643, EPI_ISL_598644, EPI_ISL_598645, EPI_ISL_598646, EPI_ISL_598647, EPI_ISL_598648, EPI_ISL_598649, EPI_ISL_598650, EPI_ISL_598651, EPI_ISL_598652, EPI_ISL_598653, EPI_ISL_598654, EPI_ISL_598655, EPI_ISL_598656, EPI_ISL_598657, EPI_ISL_598658, EPI_ISL_598659, EPI_ISL_598660, EPI_ISL_598661, EPI_ISL_598662, EPI_ISL_598663, EPI_ISL_598664, EPI_ISL_598665, EPI_ISL_598666, EPI_ISL_598668, EPI_ISL_598669, EPI_ISL_598670, EPI_ISL_598671, EPI_ISL_598672, EPI_ISL_598673, EPI_ISL_598674, EPI_ISL_598675, EPI_ISL_598676, EPI_ISL_598677, EPI_ISL_598678, EPI_ISL_598679, EPI_ISL_598680, EPI_ISL_598681, EPI_ISL_598682, EPI_ISL_598683, EPI_ISL_598684, EPI_ISL_598685, EPI_ISL_598686, EPI_ISL_598687, EPI_ISL_598688, EPI_ISL_598689, EPI_ISL_598690, EPI_ISL_598691, EPI_ISL_598692, EPI_ISL_598693, EPI_ISL_598694, EPI_ISL_598695, EPI_ISL_598696, EPI_ISL_598697, EPI_ISL_598698, EPI_ISL_598699, EPI_ISL_598700, EPI_ISL_598701, EPI_ISL_598702, EPI_ISL_598704, EPI_ISL_598705, EPI_ISL_598706, EPI_ISL_598707, EPI_ISL_598708, EPI_ISL_598709, EPI_ISL_598710, EPI_ISL_598711, EPI_ISL_598712, EPI_ISL_598714, EPI_ISL_598715, EPI_ISL_598716, EPI_ISL_598717, EPI_ISL_598718, EPI_ISL_598719, EPI_ISL_598720, EPI_ISL_598721, EPI_ISL_598722, EPI_ISL_598723, EPI_ISL_598724, EPI_ISL_598725, EPI_ISL_598726, EPI_ISL_598727, EPI_ISL_598728, EPI_ISL_598729, EPI_ISL_598730, EPI_ISL_598731, EPI_ISL_598732, EPI_ISL_598733, EPI_ISL_598734, EPI_ISL_598735, EPI_ISL_598736, EPI_ISL_598737, EPI_ISL_598738, EPI_ISL_598739, EPI_ISL_598740, EPI_ISL_598741, EPI_ISL_598742, EPI_ISL_598743, EPI_ISL_598744, EPI_ISL_598745, EPI_ISL_598747, EPI_ISL_598748, EPI_ISL_598749, EPI_ISL_598750, EPI_ISL_598752, EPI_ISL_598753, EPI_ISL_598754, EPI_ISL_598755, EPI_ISL_598756, EPI_ISL_598757, EPI_ISL_598758, EPI_ISL_598760, EPI_ISL_598761, EPI_ISL_598763, EPI_ISL_598764, EPI_ISL_598767, EPI_ISL_598768, EPI_ISL_598769, EPI_ISL_598770, EPI_ISL_598771, EPI_ISL_598772, EPI_ISL_598773, EPI_ISL_598774, EPI_ISL_598775, EPI_ISL_598776, EPI_ISL_598777, EPI_ISL_598778, EPI_ISL_598779, EPI_ISL_598780, EPI_ISL_598781, EPI_ISL_598782, EPI_ISL_598783, EPI_ISL_598784, EPI_ISL_598785, EPI_ISL_598786, EPI_ISL_598787, EPI_ISL_598788, EPI_ISL_598789, EPI_ISL_598790, EPI_ISL_598791, EPI_ISL_598792, EPI_ISL_598793, EPI_ISL_598794, EPI_ISL_598795, EPI_ISL_598796, EPI_ISL_598797, EPI_ISL_598798, EPI_ISL_598799, EPI_ISL_598800, EPI_ISL_598801, EPI_ISL_598802, EPI_ISL_598803, EPI_ISL_598805, EPI_ISL_598806, EPI_ISL_598807, EPI_ISL_598808, EPI_ISL_598809, EPI_ISL_598810, EPI_ISL_598811, EPI_ISL_598812, EPI_ISL_598813, EPI_ISL_598814, EPI_ISL_598815, EPI_ISL_598816, EPI_ISL_598818, EPI_ISL_598819, EPI_ISL_598820, EPI_ISL_598822, EPI_ISL_609219, EPI_ISL_609220, EPI_ISL_609221, EPI_ISL_609222, EPI_ISL_609223, EPI_ISL_609224, EPI_ISL_609225, EPI_ISL_609226, EPI_ISL_609227, EPI_ISL_609228, EPI_ISL_609229, EPI_ISL_609231, EPI_ISL_609232, EPI_ISL_609233, EPI_ISL_609234, EPI_ISL_609235, EPI_ISL_609236, EPI_ISL_609237, EPI_ISL_609238, EPI_ISL_609239, EPI_ISL_609240, EPI_ISL_609241, EPI_ISL_609242, EPI_ISL_609243, EPI_ISL_609244, EPI_ISL_609245, EPI_ISL_609247, EPI_ISL_609248, EPI_ISL_609249, EPI_ISL_609250, EPI_ISL_609251, EPI_ISL_609252, EPI_ISL_609253, EPI_ISL_609254, EPI_ISL_609255, EPI_ISL_609256, EPI_ISL_609257, EPI_ISL_609258, EPI_ISL_609259, EPI_ISL_609260, EPI_ISL_609261, EPI_ISL_609262, EPI_ISL_609263, EPI_ISL_609264, EPI_ISL_609265, EPI_ISL_609266, EPI_ISL_609268, EPI_ISL_609269, EPI_ISL_609270, EPI_ISL_609271, EPI_ISL_609272, EPI_ISL_609273, EPI_ISL_609274, EPI_ISL_609275, EPI_ISL_609276, EPI_ISL_609277, EPI_ISL_609278, EPI_ISL_609279, EPI_ISL_609280, EPI_ISL_609281, EPI_ISL_609282, EPI_ISL_609283, EPI_ISL_609284, EPI_ISL_609285, EPI_ISL_609287, EPI_ISL_609288, EPI_ISL_609290, EPI_ISL_609291, EPI_ISL_609292, EPI_ISL_609293, EPI_ISL_609294, EPI_ISL_609295, EPI_ISL_609296, EPI_ISL_609297, EPI_ISL_609298, EPI_ISL_609299, EPI_ISL_609300, EPI_ISL_609301, EPI_ISL_609303, EPI_ISL_609304, EPI_ISL_609305, EPI_ISL_609306, EPI_ISL_609307, EPI_ISL_629591, EPI_ISL_630322, EPI_ISL_634418 | | | |
| see above | Lighthouse Lab in Milton Keynes | Wellcome Sanger Institute for the COVID-19 Genomics UK (COG-UK) consortium | Cordelia Langford; David K. Jackson; Dominic Kwiatkowski; Ewan Harrison; Ian Johnston; John Sillitoe on behalf of the Wellcome Sanger Institute COVID-19 Surveillance Team; John Sillitoe on behalf of the Wellcome Sanger Institute COVID-19 Surveillance Team (http://www.sanger.ac.uk/covid-team); Roberto Amato; Sonia Goncalves; The Lighthouse Lab in Milton Keynes and Alex Alderton |
| EPI_ISL_595149 | Quadram Institute Bioscience | COVID-19 Genomics UK (COG-UK) Consortium | Alexander J Trotter; Alison E. Mather; Alp Aydin; Ana P. Tedim; Anastasia Kolyva; Andrew Bell; Andrew J. Page; Claire Stuart; Dave J. Baker; Gemma L. Kay; John Wain; Justin O'Grady; Leonardo de Oliveira Martins; Lizzie Meadows; Maria Diaz; Mark Webber; Muhammed Yasir; Nabil-Fareed Alikhan; Ngozi Elumogo; Nicholas M. Thomson; Rachael Stanley; Rachel Gilroy; Reenesh Prakash; Samir Dervisevic; Samuel Bloomfield; Steven Rudder; Thanh Le-Viet |
| EPI_ISL_613205, EPI_ISL_613215, EPI_ISL_613274, EPI_ISL_626788, EPI_ISL_639183, EPI_ISL_639323 | Wales Specialist Virology Centre Sequencing lab: Pathogen Genomics Unit | COVID-19 Genomics UK (COG-UK) Consortium | Alec Birchley; Alexander Adams; Amy Gaskin; Angela Marchbank; Bree Gatica-Wilcox; Catherine Moore; Jason Coombes; Joanne Watkins; Joel Southgate; Johnathan Evans; Laura Gifford; Lauren Gilbert; Lee Graham; Malorie Perry; Matthew Bull; Nicole Pacchiarini; Sally Corden; Sara Kumziene-Summerhayes; Sara Rey; Sarah Taylor; Simon Cottrell; Sophie Jones; Tom Connor |
