## Supplementary material for "Intrahost SARS-CoV-2 k-mer identification method (iSKIM) for rapid detection of mutations of concern reveals emergence of global mutation patterns": Table S5

We gratefully acknowledge the following Authors from the Originating laboratories responsible for obtaining the specimens, as well as the Submitting laboratories where the genome data were generated and shared via GISAID, on which this research is based.

All Submitters of data may be contacted directly via www.gisaid.org

Authors are sorted alphabetically.

Acknowledgement EPI_SET Identifier: EPI_SET_20220218oz

| Accession ID | Originating Laboratory | Submitting Laboratory | Authors |
| --- | --- | --- | --- |
| EPI_ISL_567538, EPI_ISL_567580, EPI_ISL_567699, EPI_ISL_567829 | Lighthouse Lab in Alderley Park | Wellcome Sanger Institute for the COVID-19 Genomics UK (COG-UK) consortium | Cordelia Langford; David K. Jackson; Dominic Kwiatkowski; Ewan Harrison; Ian Johnston; Jacquelyn Wynn; John Sillitoe on behalf of the Wellcome Sanger Institute COVID-19 Surveillance Team; Mairead Hyland; Roberto Amato; Sonia Goncalves; The Lighthouse Lab in Alderley Park and Alex Alderton |
| EPI_ISL_566610 | Lighthouse Lab in Cambridge | Wellcome Sanger Institute for the COVID-19 Genomics UK (COG-UK) consortium | Cordelia Langford; David K. Jackson; Dominic Kwiatkowski; Ewan Harrison; Ian Johnston; John Sillitoe on behalf of the Wellcome Sanger Institute COVID-19 Surveillance Team; Rob Howes; Roberto Amato; Sonia Goncalves; The Lighthouse Lab in Cambridge and Alex Alderton |
| EPI_ISL_602090 | Lighthouse Lab in Glasgow | Wellcome Sanger Institute for the COVID-19 Genomics UK (COG-UK) Consortium | Anna Dominiczak and Alex Alderton; Carol Clugston; Cordelia Langford; David Gray; David K. Jackson; Dominic Kwiatkowski; Ewan Harrison; Harper VanSteenhouse; Ian Johnston; John Sillitoe on behalf of the Wellcome Sanger Institute COVID-19 Surveillance Team; Roberto Amato; Sonia Goncalves; Yumi Kasai |
| EPI_ISL_549554, EPI_ISL_549594, EPI_ISL_549801, EPI_ISL_567311, EPI_ISL_567457, EPI_ISL_568244, EPI_ISL_568250, EPI_ISL_568273, EPI_ISL_568278, EPI_ISL_580886, EPI_ISL_580947, EPI_ISL_580977, EPI_ISL_581080, EPI_ISL_581194, EPI_ISL_588234, EPI_ISL_588727, EPI_ISL_588741, EPI_ISL_588748, EPI_ISL_588761, EPI_ISL_588805, EPI_ISL_588971, EPI_ISL_590177, EPI_ISL_590315, EPI_ISL_590670, EPI_ISL_599489, EPI_ISL_599504, EPI_ISL_599511, EPI_ISL_599561, EPI_ISL_599575, EPI_ISL_599633, EPI_ISL_599653, EPI_ISL_599686, EPI_ISL_599714, EPI_ISL_599747, EPI_ISL_600575, EPI_ISL_600587, EPI_ISL_600855, EPI_ISL_600872, EPI_ISL_601104, EPI_ISL_601108, EPI_ISL_601127, EPI_ISL_601168, EPI_ISL_601175, EPI_ISL_601206, EPI_ISL_601208, EPI_ISL_601334, EPI_ISL_601357 | | | |
| see above | Lighthouse Lab in Glasgow | Wellcome Sanger Institute for the COVID-19 Genomics UK (COG-UK) consortium | Anna Dominiczak and Alex Alderton; Carol Clugston; Cordelia Langford; David Gray; David K. Jackson; Dominic Kwiatkowski; Ewan Harrison; Harper VanSteenhouse; Ian Johnston; John Sillitoe on behalf of the Wellcome Sanger Institute COVID-19 Surveillance Team; John Sillitoe on behalf of the Wellcome Sanger Institute COVID-19 Surveillance Team (http://www.sanger.ac.uk/covid-team); Roberto Amato; Sonia Goncalves; Yumi Kasai |
| EPI_ISL_549988, EPI_ISL_550150, EPI_ISL_566436, EPI_ISL_566447, EPI_ISL_566452, EPI_ISL_566471, EPI_ISL_566484, EPI_ISL_566506, EPI_ISL_566570, EPI_ISL_566728, EPI_ISL_580915, EPI_ISL_581342, EPI_ISL_599928, EPI_ISL_599964, EPI_ISL_600016 | | | |
| see above | Lighthouse Lab in Milton Keynes | Wellcome Sanger Institute for the COVID-19 Genomics UK (COG-UK) consortium | Cordelia Langford; David K. Jackson; Dominic Kwiatkowski; Ewan Harrison; Ian Johnston; John Sillitoe on behalf of the Wellcome Sanger Institute COVID-19 Surveillance Team; John Sillitoe on behalf of the Wellcome Sanger Institute COVID-19 Surveillance Team (http://www.sanger.ac.uk/covid-team); Roberto Amato; Sonia Goncalves; The Lighthouse Lab in Milton Keynes and Alex Alderton |
