## Supplementary material for "Intrahost SARS-CoV-2 k-mer identification method (iSKIM) for rapid detection of mutations of concern reveals emergence of global mutation patterns": Table S6

### SUPPLEMENTAL TABLE

**Data Availability**

| | |
|---|---|
| GISAID Identifier: | EPI_SET_20220603uy |
| DOI: | https://doi.org/10.55876/gis8.220603uy |

All genome sequences and associated metadata in this dataset are published in GISAID's EpiCoV database. To view the contributors of each individual sequence with details such as accession number, Virus name, Collection date, Originating Lab and Submitting Lab and the list of Authors, visit https://doi.org/10.55876/gis8.220603uy

**Data Snapshot**

- EPI_SET_20220603uy is composed of 3,242 individual genome sequences.
- The collection dates range from 2019-12-26 to 2022-01-07;
- Data were collected in 192 countries and territories;
- All sequences in this dataset are compared relative to hCoV-19/Wuhan/WIV04/2019 (WIV04), the official reference sequence employed by GISAID (EPI_ISL_402124). Learn more at https://gisaid.org/WIV04.
